# DGIdb 3.0: a redesign and expansion of the drug-gene interaction database

**DOI:** 10.1101/200527

**Authors:** Kelsy C. Cotto, Alex H. Wagner, Yang-Yang Feng, Susanna Kiwala, Adam C. Coffman, Greg Spies, Alex Wollam, Nick Spies, Obi L. Griffith, Malachi Griffith

## Abstract

The Drug-Gene Interaction Database (DGIdb, www.dgidb.org) consolidates, organizes, and presents drug-gene interactions and gene druggability information from papers, databases, and web resources. DGIdb normalizes content from more than thirty disparate sources and allows for user-friendly advanced browsing, searching and filtering for ease of access through an intuitive web user interface, application programming interface (API), and public cloud-based server image. DGIdb v3.0 represents a major update of the database. Nine of the previously included twenty-eight sources were updated. Six new resources were added, bringing the total number of sources to thirty-three. These updates and additions of sources have cumulatively resulted in 56,309 interaction claims. This has also substantially expanded the comprehensive catalogue of druggable genes and antineoplastic drug-gene interactions included in the DGIdb. Along with these content updates, v3.0 has received a major overhaul of its codebase, including an updated user interface, preset interaction search filters, consolidation of interaction information into interaction groups, greatly improved search response times, and upgrading the underlying web application framework. In addition, the expanded API features new endpoints which allow users to extract more detailed information about queried drugs, genes, and drug-gene interactions, including listings of PubMed IDs (PMIDs), interaction type, and other interaction metadata.

## INTRODUCTION

The Drug-Gene Interaction database (DGIdb, www.dgidb.org) was first released in 2013 to consolidate drug-gene interactions and potentially druggable genes into a single resource with a powerful interface to query these data (1). DGIdb 2.0 was released in 2016 and featured substantial content updates, a more intuitive user interface, and the inclusion of an API (2). Despite the success of the DGIdb, 2.0 became outdated in several ways. New sources have become available since 2.0’s release and many existing sources within the database required updates. The notion of grouping *drug claims* (an assertion of a drug concept by a constituent resource) to match a canonical drug source (PubChem compounds), and grouping *gene claims* (an assertion of a gene concept by a constituent resource) to a canonical gene source (NCBI Entrez Gene) was described in the 2.0 paper (3, 4). In brief, grouping allows the DGIdb to relate disparate representations of a gene or drug concept through a core entity, or *group*. As the database grew, the grouping success rates dropped, and new classes of drugs were not fully represented in the search results. Additionally, while the DGIdb grouped drug and gene claims, interactions were still reported at the claim level, with no concept of interaction groups. Furthermore, the initial API that was released alongside 2.0 had limited endpoints that prevented users from extracting additional information about their queried drug-gene interactions, such as PubMed IDs (PMIDs) supporting the interactions and other interaction metadata. Finally, the DGIdb has grown enormously since its initial publication in 2013, and as a result, interaction searches had become significantly slower. This new release addresses these limitations, providing an improved user interface, quicker response times for searches, and new methods for accessing and incorporating the DGIdb into bioinformatics workflows.

## EXPANDED CONTENT

For 3.0, there has been a substantial expansion of the content of the database through the addition of new sources and the updating of existing sources. Six new sources have been added, which brings the total number of sources represented by the DGIdb to thirty-three. Three of these new sources provide drug-gene interactions from prominent expert-curated databases of clinically actionable variants similar to the CIViC and Clearity Foundation sources, already included in the DGIdb (Supplementary Table 1) (5). These include the Precision Oncology Knowledge Base (OncoKB), Cancer Genome Interpreter (CGI) (https://www.cancergenomeinterpreter.org/), and The Jackson Laboratory Clinical Knowledgebase (CKB) (6, 7). From these three new sources, 2,822 new interaction claims were added. These sources were added as antineoplastic interaction sources, resulting in an 36.7% increase in antineoplastic drugs. Drug-gene interactions were also added from the FDA Pharmacogenomics website (https://www.fda.gov/Drugs/ScienceResearch/ResearchAreas/Pharmacogenetics/) and the National Cancer Institute (NCI) Cancer Gene Index (https://wiki.nci.nih.gov/display/cageneindex), resulting in an additional 276 and 6,231 interaction claims, respectively. In addition to these sources, the existing druggable gene category sources were expanded with data extracted from a new druggable genome paper that used computational approaches to identify druggable genes from genome-wide association studies (Supplementary Table 2) (8). The inclusion of this source into the DGIdb results in an additional 2,300 potentially druggable genes, a 58% increase from 2.0 (Figure 1). Notably, this independent definition of the druggable genome almost completely encapsulates many other gene categories (e.g. kinases, G-protein coupled receptors) that are expected to be good drug targets (Figure 2).

**Figure 1:**
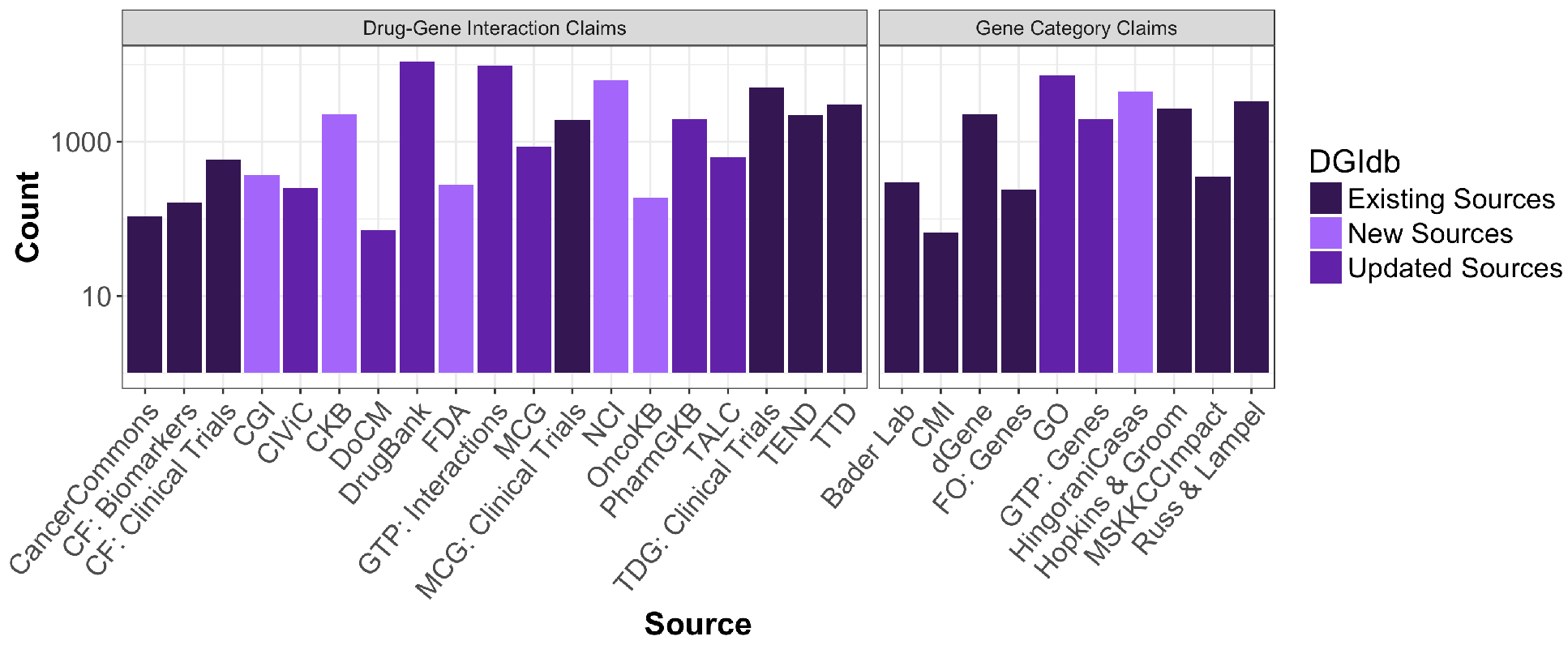
DGIdb 3.0 content by source. The number of drug-gene interaction claims (first panel) and druggable gene categories (second panel) are separated into three categories: sources that existed in the DGIdb previously, sources that existed in the DGIdb previously but have been updated for 3.0, or sources that are entirely new to the DGIdb. In total, there are currently 56,309 (18,493 new) drug-gene interaction claims, and 23,173 (4,673 new) gene category claims. Abbreviations: CF = Clearity Foundation, CGI = Cancer Genome Interpreter, CKB = JAX-Clinical Knowledgebase, CMI = Caris Molecular Intelligence, FO = Foundation One, GTP = Guide to Pharmacology, MCG = My Cancer Genome, OncoKB = Precision Oncology Knowledge Base, TALC = Targeted Agents in Lung Cancer, TTD = Therapeutic Target Database, TEND = Trends in the Exploration of Novel Drug targets, GO = Gene Ontology, and MSKCC = Memorial Sloan Kettering Cancer Center.

**Figure 2:**
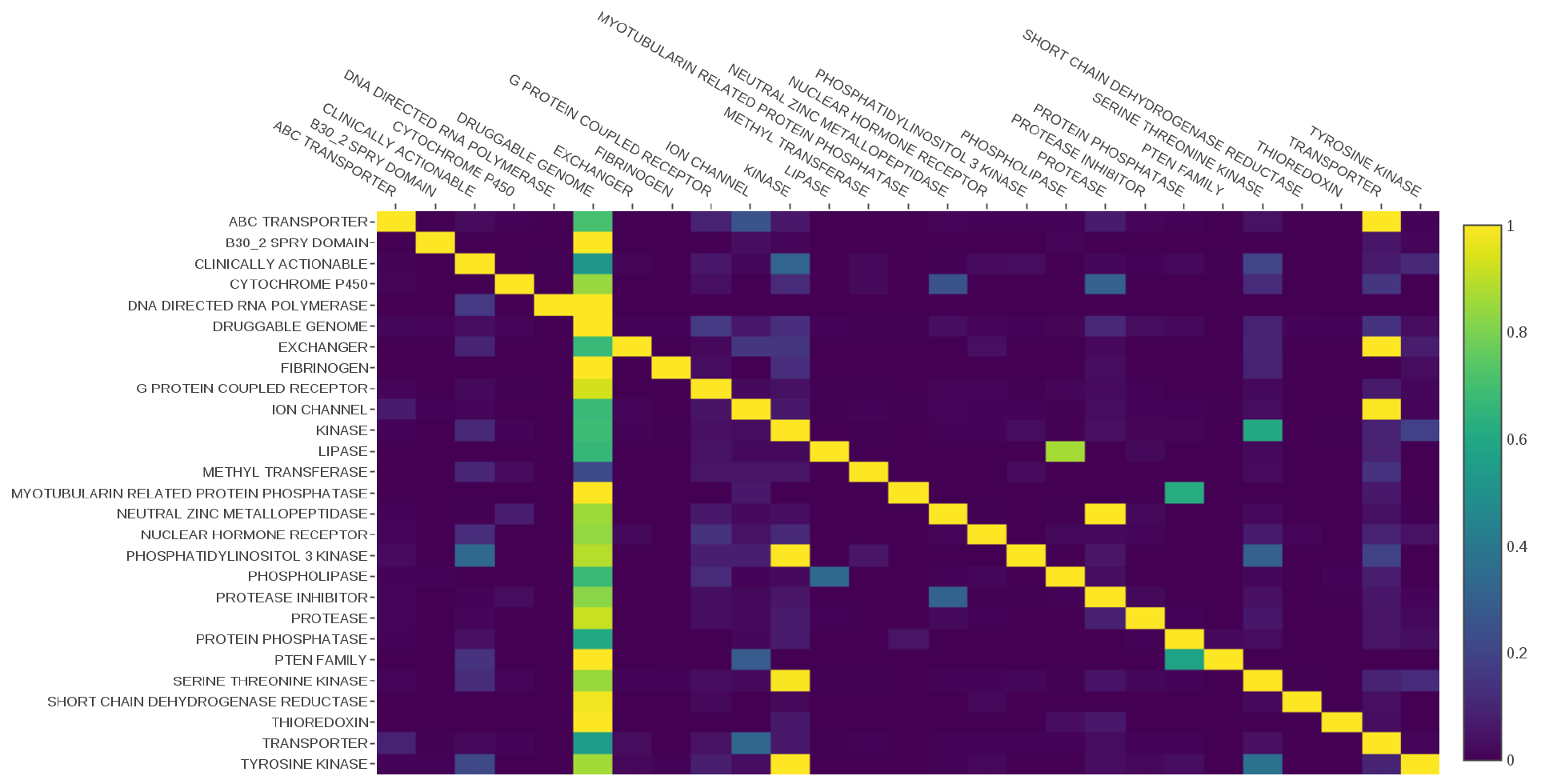
DGIdb 3.0 Druggable Genome Categories. This figure shows the percent overlap of the various gene categories of the DGIdb 3.0, with the overlap expressed as a percentage of the category along the Y-axis. For example, the Druggable Genome row has mostly low values due to only a small percentage of the genes within it matching each of the smaller gene categories (e.g. 12% of Druggable Genome genes in the Kinase category). However, the Druggable Genome column has mostly high values due to most of the categories along the y axis being near-subsets of the Druggable Genome category (68% of Kinase genes are in the Druggable Genome category).

As the DGIdb has matured, maintaining current representations of existing sources has become increasingly important relative to identifying new interaction sources. Curated updates of drug-gene interaction data from My Cancer Genome and the targeted and biologic therapies for non-small-cell lung cancer (TALC) have occurred, resulting in moderate increases in drug-gene interactions from these sources (9, 10). To help prevent regularly updated sources from becoming outdated in the DGIdb, we have rewritten and expanded *online updaters* to keep frequently updated sources current within the database. With these more frequent, incremental updates to the DGIdb, there has been a substantial increase in drug-gene interaction content from IUPHAR’s Guide to Pharmacology and CIViC, a 23% and 200% increase, respectively (5, 11). Additionally, a major update of DrugBank from version 4.3 to 5.0 was performed, resulting in a 30% increase in interaction claims from this source (12). Gene Ontology was also updated which provided a moderate increase for the druggable gene categories (13). For this release, 18,493 new interaction claims were added, of which 50% were from updated sources and 50% from new sources. In total, there are now 6,106 druggable genes and 29,783 drug-gene interactions, which cover 41,100 genes and 9,495 drugs, within the DGIdb.

The enhancements to the online updaters have also been applied to Entrez Gene, from which, 99% of all gene claims made by the DGIdb constituent sources were grouped (Supplemental Figure 1) (3). Another major change from 2.0 to 3.0 is that the canonical drug source for the DGIdb has switched from using PubChem compounds to ChEMBL molecules (14). This switch has added 1.7 million ChEMBL molecules to the database for potential matching to drug claims. Importantly, switching to ChEMBL has added 195 antibody drugs (e.g., trastuzumab, cetuximab), a drug class that is absent from the PubChem database and frequently requested by users. These antibody drugs matched to 539 distinct drug claims from the constituent sources of the DGIdb. With ChEMBL as the canonical drug source and the improvements to the grouping strategy below, 80.2% of all drug claims now group. Many of the resources we pull from strive to be as comprehensive as possible, and sometimes include broad classes of drug or therapy (e.g. “hormone therapy”, “mtor inhibitors”, “chemotherapy”, “radiation”, “antibiotics”, etc.), which account for a large percentage of the remaining drug claims.

## NEW FEATURES AND ENHANCEMENTS

A major feature added in 3.0 is a user-selectable series of preset filters. These allow users to select drug-gene interactions that align with some of the most common search use cases. While it was possible for a user to filter on some of the following attributes in 2.0, these concepts have now been structured as stand-alone filters that can help focus a user’s search results with a single click. The filters currently include FDA approved drugs, antineoplastic drugs, immunotherapy drugs, clinically actionable genes, genes included in the druggable genome definition, and drug resistant genes. *FDA approval status* is extracted from ChEMBL v23 (current as of this writing) (14). *Antineoplastic drugs* are defined by inclusion in an antineoplastic drug-gene interaction source (e.g. My Cancer Genome), or as a drug with an antineoplastic attribute from its constituent source. Immunotherapy drugs are defined as any drug with an attribute of “immunosuppressive agent”, “immunomodulatory agent”, or “immunostimulant”. *Clinically actionable* genes are genes that constitute the DGIdb “clinically actionable” gene category, and by definition is used to inform clinical action (e.g. the Foundation One diagnostic gene panels). Similarly, *druggable genome* genes are genes listed in the DGIdb “druggable genome” gene category (8, 11, 13, 15–19). *Drug resistance* genes are defined by the Gene Ontology as genes that confer drug resistance or susceptibility (GO identifier 0042493), and are maintained in the DGIdb through the “drug resistance” gene category (13). To incorporate these new filters, we have redesigned the data model, expanded the definition of the druggable genome, and restructured the UI to include a redesigned search form and results interface. The UI also now features drug, gene, and interaction views to leverage the new grouping strategies and preset filters.

Since the last release, several changes have been made to the drug grouping strategy to improve overall grouping percentages. We have added support for fuzzy searching when grouping if direct matches to drug groups are not found, enabling grouping of drug claims with slightly different means of joining multiword terms (whitespace, dashes, underscores). Code has also been added to identify and remove aliases that are highly ambiguous to improve exact matching of drug claims.

In addition to improved drug grouping, we have added interaction grouping, linking together interaction claims from multiple sources that describe the exact same drug-gene interaction. The grouping efficiency for interactions is 75.2%. The successful grouping of an interaction claim is dependent on the associated gene claims and drug claims successfully grouping. As a result, interaction grouping percentages are closely related to the grouping percentages for drugs and genes (Supplemental Figure 1). The introduction of interaction groups has led to a noticeable improvement in response times on the UI for interaction searches, even when query sizes and result sizes grow larger (Figure 2). Before interaction groups were created, searches in 2.0 had to query almost twice as many database tables as in 3.0. Interaction groups allow for more efficient queries, which leads to a 14-fold reduction in response times when searching the DGIdb.

We have expanded and added additional API endpoints that now allow users to extract more information on the drug-gene interactions provided by the DGIdb. These include endpoints to list all the interaction groups, gene groups, and drug groups in the DGIdb, as well as endpoints to view detailed information about an individual interaction, gene, or drug group. Drug/gene/interaction group endpoints include various metadata about the group including its constituent claims. The information presented in the interaction search results endpoint has been expanded to include the Entrez and Chembl ID of the gene and drug involved in an interaction, as well as a list of publications that support each interaction. Users can also apply the preset filters to their interaction search through the API. This brings the interaction search endpoint in sync with the user interface. Moreover, the information presented in the interaction search results and the new individual endpoints for interactions, genes, and drugs more closely mirrors the views available via the user interface. These new endpoints allow users to more efficiently export all of the data available in the DGIdb.

## USER INTERFACE (UI) UPDATES

The 3.0 release of the DGIdb features a dramatic overhaul of the UI that reflects many of the backend changes detailed above. The search interactions page now allows the user to apply the new preset filters in addition to the existing advanced filters (e.g. Source Databases, Gene Categories, and Interaction Types) directly on the search form (Figure 4). On performing a search, the user is redirected to the redesigned search interactions results page, enabled by a new concept of interaction groups instead of individual interaction claims. This new view displays search results as lists of visually distinct panels for each search term mapped to gene or drug groups. A list of uniquely matched terms is shown in one tab whereas a second tab summarizes the list of ambiguously matched search terms and unmatched terms. Each panel displays a table of interaction groups, along with summaries of the respective interaction types, sources, supporting publication PMIDs, and a ranking. Users interested in a particular interaction can navigate from the interaction search results to the corresponding interaction group view. Similarly, detailed information about an interacting gene or drug can be found in the associated gene or drug group view (Supplemental Figure 2). These new group views feature (1) a *summary tab*, which details all relevant information collected from the various source claims (e.g. aliases, FDA approval status, supporting publications); (2) an *interactions tab* (for gene and drug group views), which lists summary panels of each interaction in the DGIdb for this gene or drug; and (3) a *claims tab* with detailed information about each claim supporting the group.

## USAGE AND ACCESSIBILITY

The utility of the DGIdb as a resource is reflected in its substantial web and especially API traffic. The website receives ~1,700 unique users and ~2,800 sessions a month, an increase of 42% and 39%, respectively, since the initial release of the DGIdb. Additionally, 42% of unique visitors within the past year were returning users. Within the last year, the DGIdb API has been used by 3,689 unique IP addresses for a total of 939,429 requests, as seen in Supplemental Figure 3. At the time of publication, 2.0 was used by a number of bioinformatics tools in the development of their platforms. DGIdb has since been integrated into four additional tools, bringing the total number of known platforms that utilize the DGIdb to ten. These platforms are GeneCards (http://www.genecards.org), CVE (20), rDGIdb (21), PANDA {Hart, 2015}, iCAGES (icages.usc.edu), BioGPS (22), Omics Pipe (23), GEMINI (24), StationX (www.stationxinc.com), and IHLDB.rf (www.lungcancerdatabase.com). These integrations highlight the accessibility and utility of the DGIdb API, which has been expanded to include new endpoints in 3.0.

**Figure 3:**
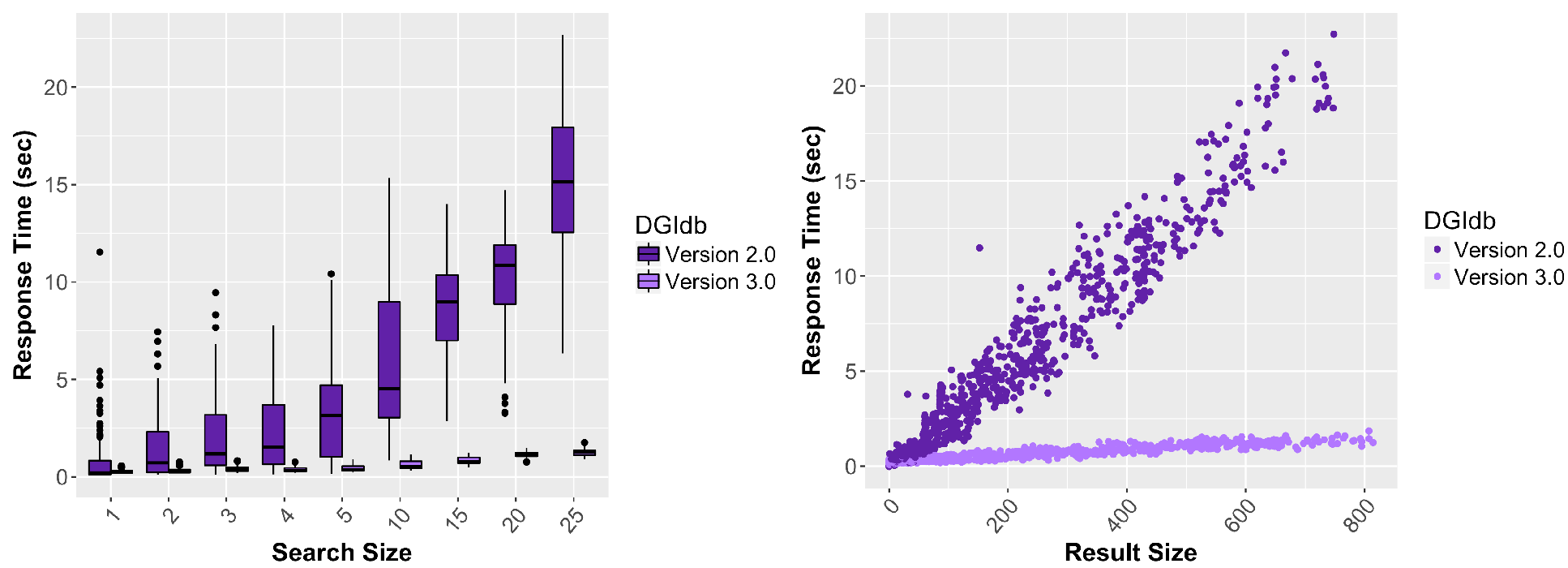
Comparison of the DGIdb 2.0 and 3.0 response time by search and result size. The first panel summarizes observed response times (n=100) for searches of various sizes (1, 2, 3, 4, 5, 10, 15, 20, and 25) of randomly selected genes. The second panel summarizes observed response times for the corresponding interaction search results. Note that the size of interaction results varied widely (range: 0 to 814) depending on the size and gene identities of the search. Response times were dramatically reduced in DGIdb 3.0 across all search and result sizes tested/observed.

**Figure 4:**
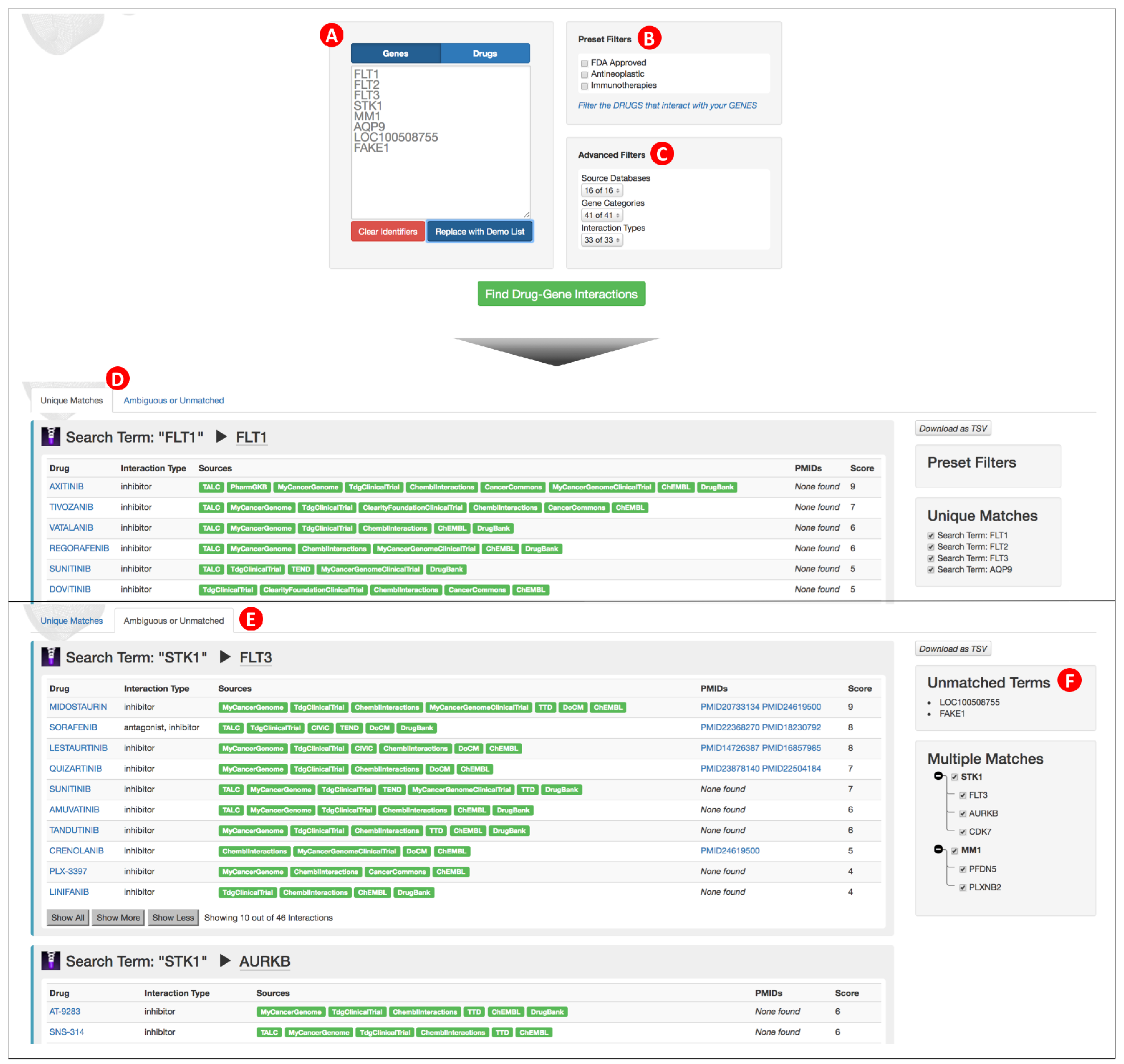
The DGIdb interaction search interface. (**A**) A search field accepts drug or gene identifiers, depending on which tab is selected, and provides autocompletion suggestions for search terms. (**B**) Preset options are provided to filter search results based on the attributes of mapped interactors. Preset filter options vary based on the type of query supplied (genes vs. drugs) and do not apply to the queries themselves. (**C**) Advanced filters allow the user to further filter based on source database, gene category, or interaction type. (**D**) The interaction search results (Unique Matches) view shows results for search terms that were matched within the DGIdb. This view provides a summary of linked genes and drugs with information about which sources along with any PubMed IDs (PMIDs) that reported this interaction. (**E**) The interaction search results (Ambiguous or Unmatched) view shows search terms that were either ambiguously matched or unmatched within the DGIdb. This view provides the same information that is reported for terms that were matched within the DGIdb. (**F**) Additional side panels provide a brief summary of unmatched and ambiguously matched terms. For terms that are ambiguous, the user can choose to select or deselect terms to control which interaction results they see.

To handle the increasing usage of the DGIdb, the database backend has undergone significant updates. One of the most notable is the upgrade from Rails 3 to the Rails 5 framework. We also added 22 new tables to the PostgreSQL database schema consisting of 2,326,676 records supporting the described changes, and added 4,065 new lines of code to the repository, an expansion of the codebase by 18.9%.

This activity can be seen through the commit history to the GitHub repository in Supplemental Figure 4. For access to this code and data, please see the availability section below.

The open source DGIdb software was previously available under the GNU General Public License 3.0 (GPL3) and has been re-released under the more permissive MIT license. The data contained within the DGIdb are available under the licenses assigned by their host sources which makes it possible to integrate the DGIdb into any workflow. Since 2.0, the database has moved from being hosted on a local server to being hosted on Amazon Web Services (AWS). With this change in server host, the availability of the DGIdb has been expanded through the release of two Amazon Machine Images (AMIs), a development environment and production environment. By providing this production environment for users, we now support a quick-start solution for private, cloud-based applications.

## CONCLUSIONS AND FUTURE DIRECTIONS

DGIdb 3.0 has undergone significant changes. The number of drug-gene interactions and druggable genome definitions has been substantially expanded through the updating of existing sources and inclusion of new sources. To improve searches, we have included preset filters that allow searches for commonly requested gene or drug classes. To utilize these filters, a more intuitive UI was created and new fields to the database schema were added. Additionally, there are new ways to both access the data within the DGIdb and to deploy local or cloud instances of the database.

While the DGIdb remains a powerful resource for querying drug-gene interactions, we anticipate future changes that will improve the user experience and the content within it. One change is that claim information will be linked to the licensing terms from constituent sources through the various methods of accessing the DGIdb including the web interface, API endpoints, and data downloads. Secondly, while grouping statistics for claims have improved in the DGIdb 3.0, there are still a significant percentage of claims that remain ungrouped. To address this, the the drug grouper will need to be optimized to handle the exceptions that currently prevent successful grouping. Third, much of the user interface is a series of static renders by the server. A potential future enhancement is a more reactive, client-side web application that would allow for more dynamic visualization and exploration of the data. Finally, other databases (e.g. CIViC) have had success utilizing a community curation model. In an effort to enable community feedback to address the complexity of grouping and representing drug-gene interaction data, the DGIdb will be moved to a similar model. Expected changes include users creating login accounts, suggesting sources or claims for inclusion, and verification that a source or claim has enough support to be included within the DGIdb. These changes will require significant code updates but will greatly increase the utility of the DGIdb.

## AVAILABILITY

DGIdb is an open access database and web interface (www.dgidb.org) with open source code available on GitHub (https://github.com/griffithlab/dgi-db). We also provide data downloads for drug claims, gene claims, and interaction claims on the website in addition to a SQL data dump (http://dgidb.org/downloads). Information about the API and its endpoints can also be found on the website (http://dgidb.org/api).

## SUPPLEMENTARY DATA

Supplementary Data are available at NAR online.

This work is supported by the National Human Genome Research Institute [R00HG007940] to M.G.0.L.G. is supported by the National Cancer Institute [K22CA188163]. A.H.W is supported by the NCI [T32CA113275, F32CA206247]. K.C.C. is supported by National Cancer Institute [T32CA113275] and National Institute of General Medical Sciences [5R25GM103757]. The incorporation of CIViC and related interaction data sources was supported by the National Cancer Institute [U01CA209936]. Funding for open access charge: National Institutes of Health [R00HG007940].

